# Nose-to-Brain Healing: Hypoxia-Preconditioned Mesenchymal Stem Cells Prompt Recovery in Hypoxic-Ischemic Encephalopathy Rats

**DOI:** 10.1101/2025.01.21.634067

**Authors:** Inês Serrenho, Alexandre M. Carvalho, Inês Caramelo, Beatriz Araújo, Carla M.P. Cardoso, Bruno Manadas, Graça Baltazar

**Affiliations:** RISE-Health, Department of Medical Sciences, Faculty of Health Sciences, University of Beira Interior, Av. Infante D. Henrique, 6200-506 Covilhã, Portugal; CNC-UC - Center for Neuroscience and Cell Biology, University of Coimbra; CiBB - Centre for Innovative Biomedicine and Biotechnology, University of Coimbra; Crioestaminal, Stemlab S.A., Cantahede, Portugal

## Abstract

Neonatal HIE poses a significant risk factor for neurodevelopment impairment. Therapeutic hypothermia, the current standard of care for this condition, has several constraints and reduced effectivity, especially in more severe cases. Thus, it is necessary to explore novel therapeutics, like MSCs. Although previous studies report that administration of MSCs (from different sources) prompted the recovery of HIE-lesioned animals, high doses are currently used. First, this study compared the efficacy of IN versus IV administration of 50,000 UC-MSCs in a rat model of neonatal HI brain injury. For this cell dose, only IN-UC-MSC therapy reduced infarct volume, an effect accompanied by improvements of motor skills and recognition memory. Also, IN-UC-MSC administration restored myelination in the corpus callosum and mitigated glial reactivity more effectively than IV administration. In a second part of the study, to potentiate the effect of UC-MSCs administration, postnatal rats that underwent HI injury received 25,000 hypoxia-preconditioned UC-MSCs or its secretome two days later, via IN route. The administration of a low-dose of hypoxia-preconditioned UC-MSCs was sufficient to induce neurological recovery and modulation of glial response. Moreover, the administration of the secretome of these cells was enough to induce the same extent of recovery. These findings support the higher potential of IN-UC-MSC administration, compared to IV administration, while enhancing our understanding of hypoxia-preconditioning and the role of the MSC’s secretome in driving a positive therapeutic response, contributing to the development of more effective and feasible treatments for neonatal HIE.

## 1. Introduction

The current standard of care for term HIE is TH, which has demonstrated reduced efficacy in decreasing mortality and neurodevelopmental impairment of affected neonates (1). However, TH benefits are limited to specific patient populations, since it requires specialized teams and should be initiated up to 6 hours after birth. In the US, only 10.9% of HIE patients are treated with TH (2), and in the UK, TH was applied to 41–67% of the HIE patients (3, 4). Clinical trials revealed that TH effectively reduces the mortality of HIE-diagnosed neonates when compared to untreated infants (10% vs. 20–30%)(1, 3, 4), but the previously mentioned studies suggest that there is still a significant percentage of cases that do not recover after TH.

These limitations highlight the need for more effective therapeutic strategies targeting multiple molecular and cellular pathways triggered by HI brain injury. Cell therapy with MSCs has emerged as a promising approach for treating neonatal HI brain injury. MSCs possess immunomodulatory, anti-inflammatory, and neuroprotective properties, which make them attractive candidates for neurorepair. By modulating the immune response, reducing neuroinflammation, and promoting tissue regeneration (5), MSCs hold the potential to promote neural repair and improve functional outcomes in neonates with HIE. A systematic review has reported that MSCs therapy can reduce the neurological deficits caused by HI insults to the developing brain of rodents (6). Nonetheless, the studies identified in this systematic review use high doses of MSCs per kilogram of body weight, that require extensive *in vitro* expansion of MSCs. Thus, it is necessary to investigate strategies that could decrease the number of cells used while maintaining treatment efficacy. Moreover, it is becoming increasingly evident that the secretome (i.e. range of factors secreted by the MSCs) is the main driver of the therapeutic effects of MSCs in neurological conditions. The secretome is composed of soluble factors (including cytokines, growth factors, and other proteins) and extracellular vesicles, namely exosomes and microvesicles (471). On the other hand, hypoxia preconditioning was reported to alter the proteomic profile of MSCs and enhance the secretion of various trophic factors (364), including growth factors (e.g., HGF, VEGF, bFGF, TGF-β1, IGF, BDNF, GDNF) (419, 510, 511), cytokines (339, 471, 477), and matrix modulators (e.g., angiogenin, tissue inhibitor of metalloproteinase-1, and matrix metalloproteinases), which are crucial for tissue repair and angiogenesis (161, 387, 475, 508). Also, hypoxia-activated pathways, such as the ErK1/2-MAPK and PI3K/Akt pathways, regulate fibroblast migration and neurogenesis (419, 512), further enhancing tissue repair and regeneration in various injury models, including conditions affecting the central nervous system. Hypoxia exposure also significantly influenced the biogenesis, release, and content of MSC’s extracellular vesicles. Studies described that hypoxic-preconditioning increased the expression of various exosomal microRNAs, such as miR-21-5p, which has been found to regulate cellular processes like glial polarization while having an antiapoptotic effect (513). Chapter 5 also supports the use of hypoxia as a strategy to enhance the therapeutic potential of UC-MSCs in a HIE rodent model, with animals treated with the preconditioned cells presenting improved neurological outcomes and modulation of brain pathways that were dysregulated in HIE rats.

In this context, the objective of the present study is to optimize stem cell therapy for neonatal HIE, by adjusting culture and administration procedures that allow for the reduction of the dose of UC-MSCs necessary to induce neurological recovery. First, we evaluated the impact of the UC-MSCs administration route on brain damage and neurodevelopmental outcomes in a neonatal rat model of HI brain injury. Second, we assessed if IN-delivered hypoxia-preconditioned UC-MSCs could cumulatively increase the efficacy of these strategies. Finally, we investigated if IN-delivery of the secretome obtained from the hypoxia-preconditioned UC-MSCs had the same potential for neurological recovery in this model.

## 2. Results

### 2.1. Intranasal administration of UC-MSCs decreased neurological injury in neonatal rats more efficiently

To evaluate the impact of the UC-MSCs administration route on neonatal brain damage and neurodevelopmental outcomes, using low doses of MSCs, a neonatal HI brain lesion was induced in P10 Wistar rats, and, two days later, the animals were treated with low doses of UC-MSCs delivered intravenously, HIE+MSC(IV) group, or intranasally, HIE+MSC(IN) group. Brain lesion extension was assessed 30 days after injury (Figure 1). The results show that contrary to IV-delivered UC-MSCs, IN administration of UC-MSCs significantly reduced HI brain lesion to an extent that was not significantly different from the control group.

**Figure 1.**
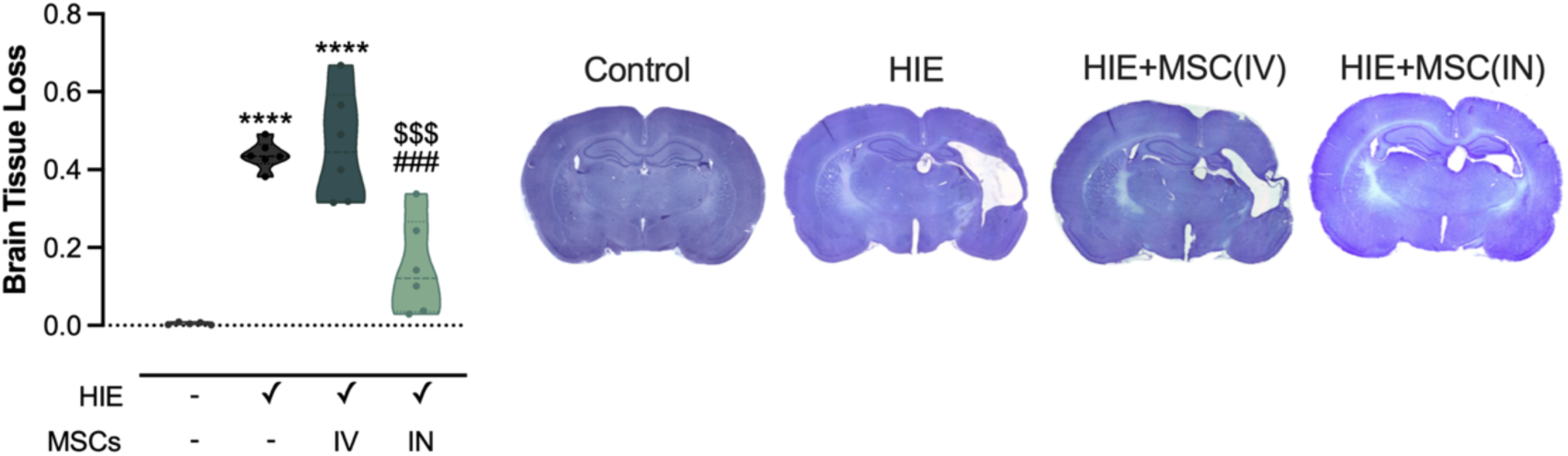
Impact of UC-MSCs administration on HI brain lesion extension. Brain lesion extension at P40 was quantified with the QuPath software using brain sections stained with Cresyl Violet. Images were acquired in a widefield microscope with a robotic stage using a 5× objective (**** p < 0.0001 vs. control, ### p < 0.001 vs. HIE, $$$ p < 0.001 vs. HIE+MSC(IV), details in Table S1).

Since brain lesion extension was considerably decreased in lesioned animals treated with IN administration of UC-MSCs, it was evaluated if the reduction was correlated with improvements of motor and cognitive functions.

The Negative Geotaxis Reflex test was used to evaluate motor coordination and vestibular reflexes of neonatal rats. Rats were placed on a 45° plane, and the time taken to turn 180° and face upwards was measured. At P14, the HIE+MSC(IN) group exhibited an improvement in the time taken to complete this test compared to the HIE group and the HIE+MSC(IV) group (Figure 2A). Similar improvements were observed at P17, with the HIE+UC-MSC(IN) group compared to the HIE group (Figure 2B). The time taken by the animals from HIE+MSC(IN) group to perform the test at both time points did not differ from the control group, suggesting the development of motor coordination and balance similar to non-lesioned animals.

**Figure 2.**
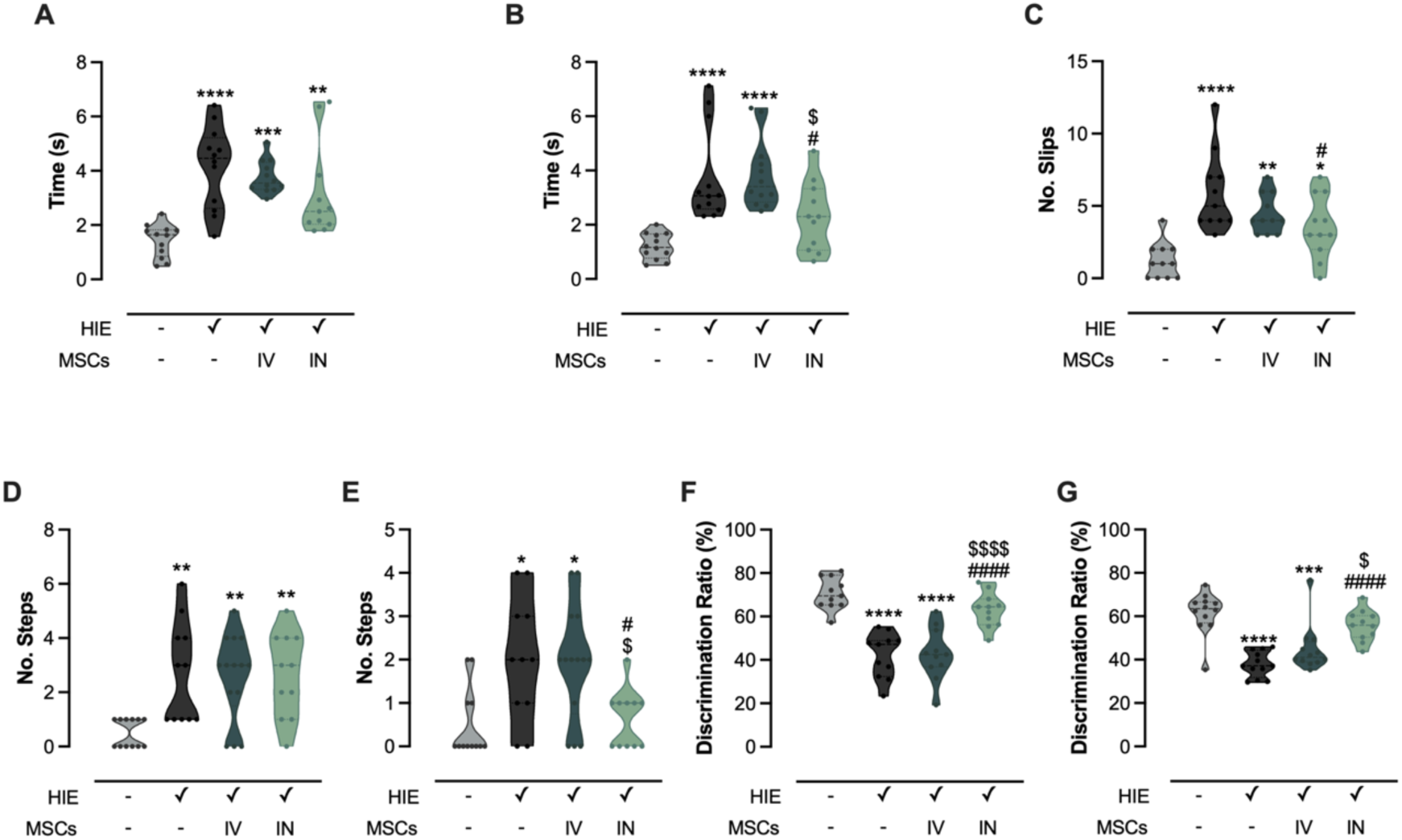
Impact of the administration route on functional outcomes in neonatal HI-injured animals. (A, B) Negative geotaxis reflex assessed at P14 and P17 by measuring the time (seconds) required to complete the task. (C) Ladder walking test performed at P30, with the number of slips recorded during the trial. (D, E) Footprint test at P28 evaluating the number of steps with overlapping and dragging. (F, G) Recognition memory evaluated at P21 and P38, presented as the discrimination ratio. Statistical significance is indicated as follows: p < 0.05, ** p < 0.01, *** p < 0.001, **** p < 0.0001 vs. control; p < 0.05, #### p < 0.0001 vs. HIE; $ p < 0.05, $$$$ p < 0.0001 vs. HIE+MSC (IV). Details are provided in Table S1.

Motor coordination and balance was evaluated by measuring the number of slips made in the Ladder Rung Walking Test (Figure 2C). Only animals that were treated with IN administration of UC-MSCs showed recovery when compared with untreated lesioned animals. Stride abnormalities were also evaluated, namely overlapping (Figure 2D) and dragging steps (Figure 2E) with the footprint test. Concerning overlapping steps, none of the therapeutic approaches counteracted the increase observed in lesioned animals. However, when analyzing the dragging steps, which are associated with a lack of motor strength, improvements were observed in the HIE+MSC(IN) group compared to the HIE group. Moreover, this parameter was ameliorated in animals treated with intranasal administration of UC-MSCs when compared to those treated with intravenous administration. The Novel Object Recognition test assessed the memory and recognition ability of the animals at two different stages of development, P21 (Figure 2F) and P38 (Figure 2G). At P21, both the HIE and HIE+UC-MSC(IV) groups presented deficits compared to the control group. However, IN-delivery of UC-MSCs led to an increase in the discrimination ratio compared to both HIE and HIE+UC-MSC(IV) groups, with no difference observed when compared to the control group. The results obtained at P38 showed a similar pattern.

These results suggest that IN-delivery of 50,000 UC-MSCs led to improved motor function and recognition memory when compared with untreated- and IV-treated lesioned animals. Additionally, this improvement is sustained in the long term. The IV administration of UC-MSCs required higher cell doses to trigger protective effects on motor and cognitive outcomes (Supplementary Figure 1).

### 2.2. Intranasally-administered UC-MSCs reduced HI-induced glial reactivity and white matter injury in the corpus callosum

As shown by previous findings, neonatal HI injury triggers a neuroinflammatory response accompanied by prolonged activation of glial cells, particularly astrocytes and microglia (7). To assess the effect of IN or IV delivery of UC-MSCs on glial reactivity, astrocytes (GFAP^+^ cells) and microglia (Iba1^+^ cells) were labeled on brain sections correspondent to the peri-infarct region (Figure 3A).

**Figure 3.**
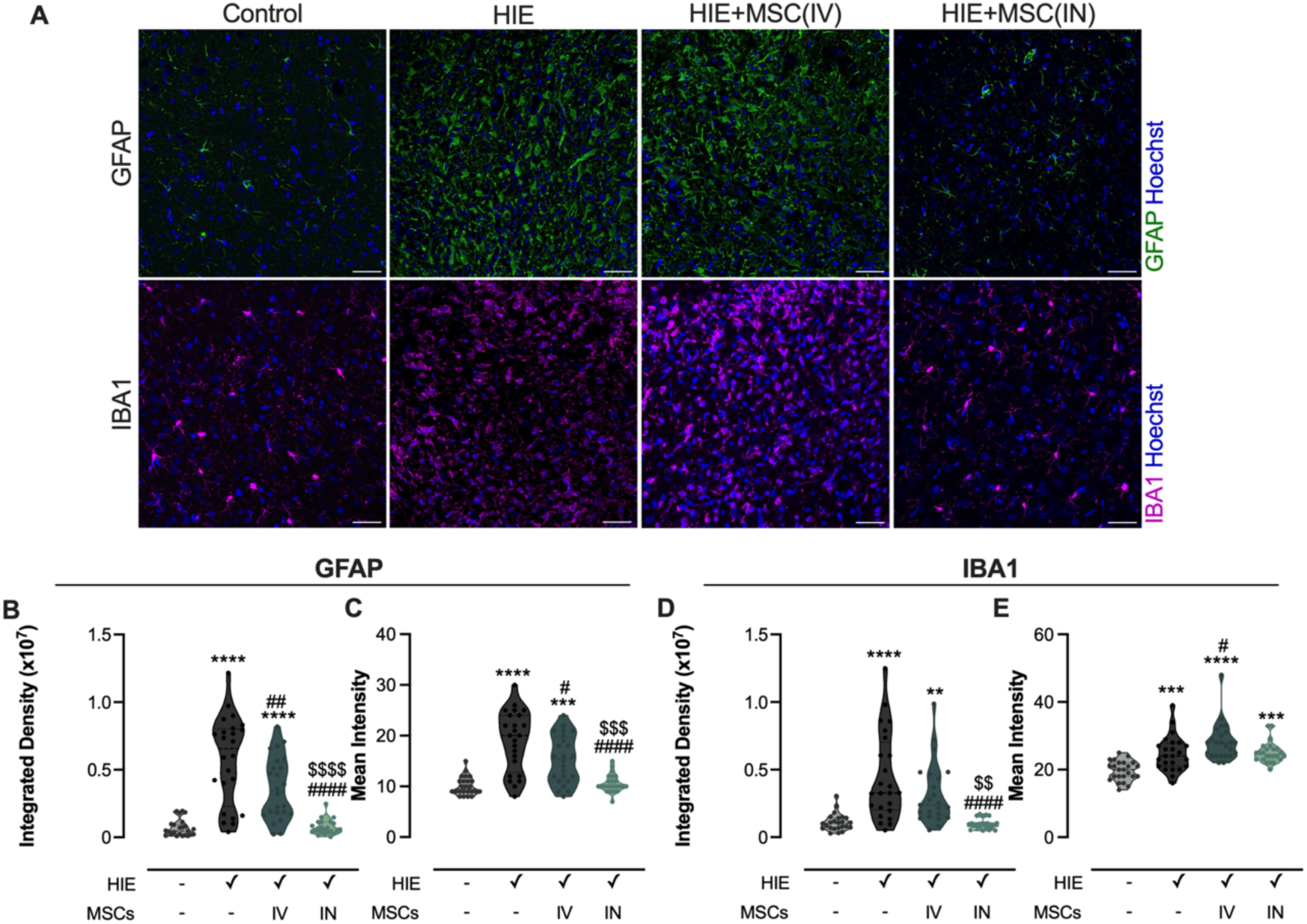
Impact of UC-MSCs administration on astrocytes and microglia 30 days after neonatal HI brain injury. (A) Representative brain sections from P40 Wistar rats immunolabelled for GFAP (green), Iba1 (magenta), with nuclei stained with Hoechst 33342 (blue). Images were acquired with a 20× objective (scale bar = 45 µm). (B, C) GFAP integrated density and mean intensity, respectively. (D, E) Iba1 integrated density and mean intensity, respectively. Statistical significance is represented as follows: ** p < 0.01, *** p < 0.001, **** p < 0.0001 vs. control; # p < 0.05, ## p < 0.01, ### p < 0.001, #### p < 0.0001 vs. HIE; $$ p < 0.01, $$$ p < 0.001, $$$$ p < 0.0001 vs. HIE+MSC(IV). Details are provided in Table S1.

The HIE group presented increased GFAP integrated and mean intensity (Figure 3B and C) in the peri-infarct area, indicators of astrogliosis and proliferation or recruitment of astrocytes to this region. UC-MSCs, administrated either IV or IN, reduced GFAP integrated density and mean intensity, with IN administration of UC-MSCs reducing GFAP labeling to control levels. These results suggest that UC-MSCs administration reduces astrocyte reactivity in the HIE model, being the IN administration more effective in mediating this effect. Microglia reactivity was also assessed (Figure 3D and E) by quantifying Iba1 levels in the peri-infarct brain region. Lesioned animals and those treated with IV-delivered UC-MSCs presented increased Iba1 integrated density and mean intensity, when compared with the control group, compatible with microglia recruitment or proliferation to this region, suggestive of increased microglial reactivity. IN-delivered UC-MSCs were able to decrease Iba1 integrated density to control levels but not Iba1 mean intensity.

Regarding microglia morphology, 3D morphological analysis of Iba1 staining in the perilesional area showed distinct microglial morphologies among the experimental groups (Figure 4A). Control and HIE+MSC(IN) animals presented a more ramified morphology, while microglia from the HIE and HIE+MSC(IV) groups displayed an ameboid shape (Figure 4A).

**Figure 4.**
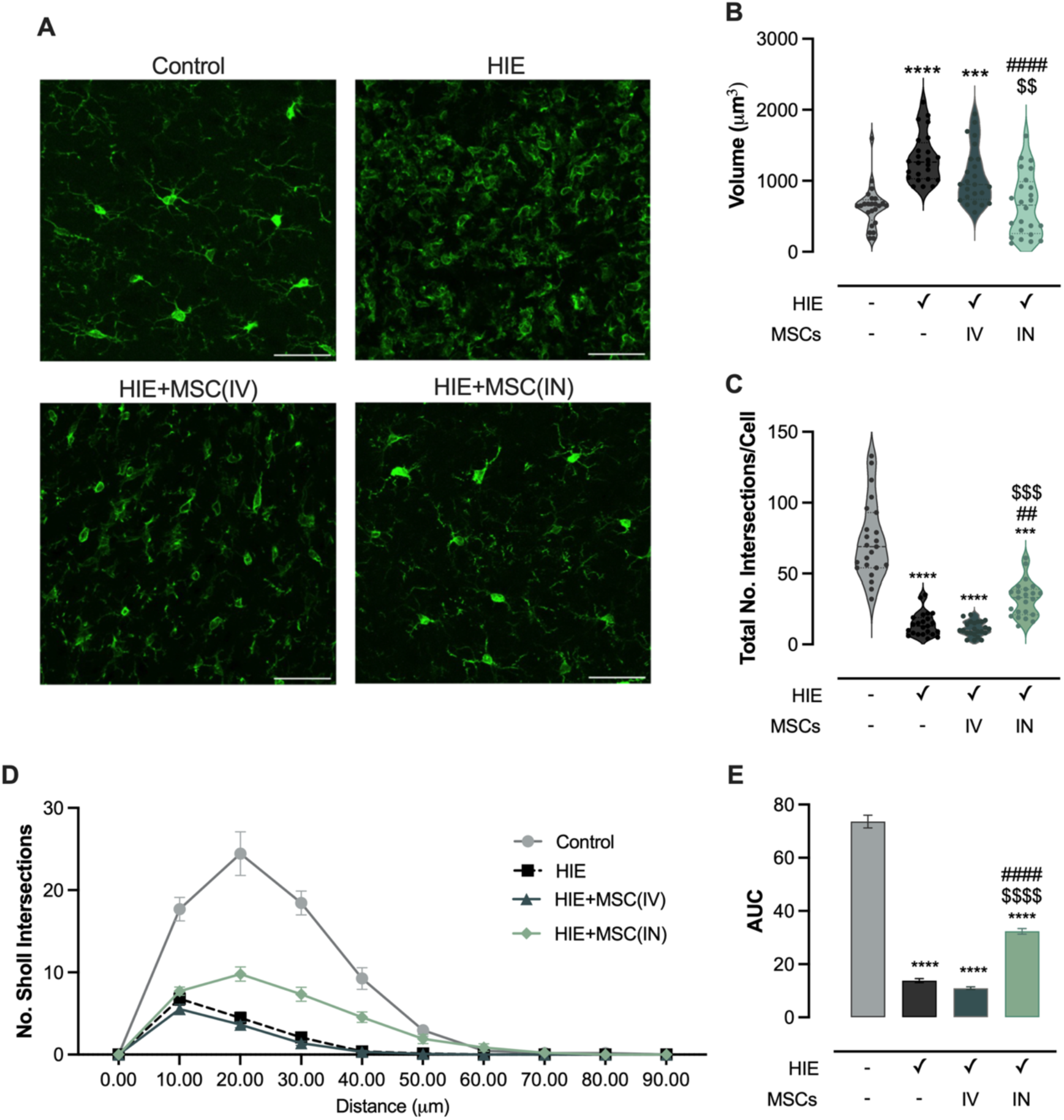
Changes in microglia morphology 30 days induced by UC-MSCs administration. (A) Representative brain sections from P40 Wistar rats labeled for Iba1. 3D images were captured using a 20× objective with the z-stack function (scale bar = 45 µm). (B) Quantification of Iba1^+^-cell volume. (C-E) Sholl analysis presented as total number of intersections, number of intersections over distance from soma and area under the curve (AUC), respectively. Statistical significance is indicated as: *** p < 0.001, **** p < 0.0001 vs. control; ## p <0.01, #### p < 0.0001 vs. HIE; $$ p < 0.01, $$$ p < 0.001, $$$$ p < 0.0001 vs. HIE+MSC(IV). Details are provided in Table S1

The microglia morphological changes translated into an increase in cell volume in HI injured animals (Figure 4B) and a significantly lower volume of the HIE+MSC(IN) group compared with the group treated with IV-delivered UC-MSCs (Figure 4B). Evaluation of branching complexity with Sholl analysis revealed a reduction in the number of intersections in the perilesional area in all HIE groups (UC-MSC-treated and untreated), indicative of decreased branching (Figure 4C). Additionally, compared to control, these groups presented fewer intersections at higher radial distances from the soma (Figure 4D and E). When compared to the HIE group, only the treatment with IN-delivered UC-MSCs partially restored microglia branching (Figure 4C, D and E).

White matter injury is a prevalent consequence of perinatal hypoxia-ischemia. Thirty days post-HI insult, the corpus callosum exhibited reduced thickness in lesioned animals (Figure 5A), accompanied by the decreased mean intensity of MBP labelling (Figure 5B) and a decline in the MBP-positive area (Figure 5C), indicating a reduction in myelination.

**Figure 5.**
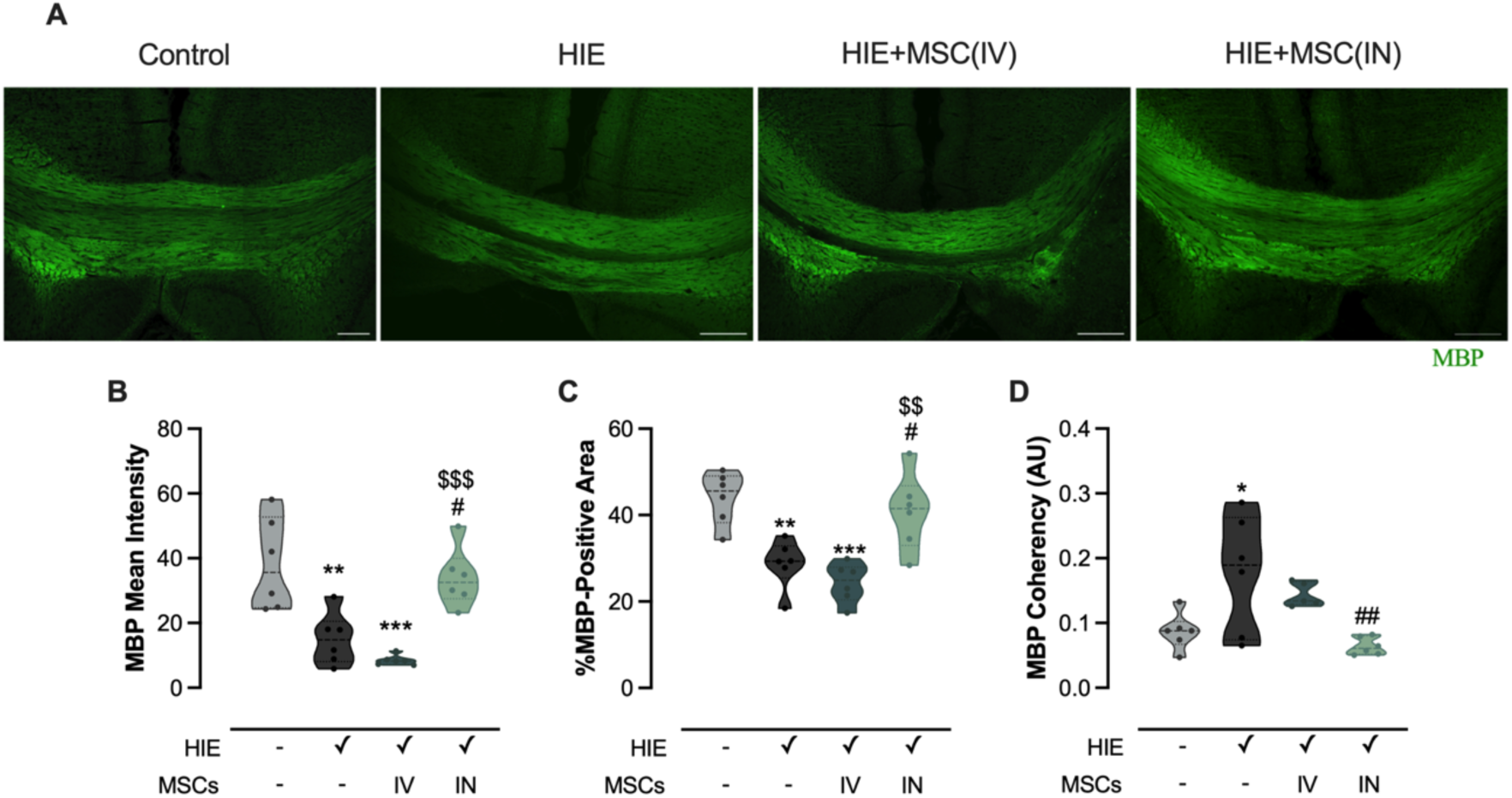
Impact of IV- and IN-UC-MSCs administration on myelin in the corpus callosum. (A) Representative brain sections from P40 Wistar rats labelled for MBP. Images were acquired with a 20× objective using the tile function (scale bar = 250 µm). (B, C and D) Quantification of MBP mean intensity, percentage of MBP-positive area, and MBP coherency, respectively. Statistical significance is indicated as: * p < 0.05, ** p < 0.01, *** p < 0.001 vs. control; # p < 0.05, ## p < 0.01 vs. HIE; $$ p < 0.01, $$$ p < 0.001 vs. HIE+MSC(IV). Details are provided in Table S1.

The coherency of MBP staining, which is inversely related to myelination complexity (8), was increased in HIE-lesioned animals compared to controls (Figure 5D), corroborating earlier findings. Notably, IN administration of UC-MSCs increased both the intensity and the area of MBP labelling more effectively than the IV administration of UC-MSCs (Figure 5B and C). Additionally, UC-MSCs IN-delivered augmented myelination complexity compared to untreated animals (Figure 5D).

### 2.3. Intranasal administration of SSH-UC-MSCs or its secretome improves neurofunctional recovery and reduces HI injury

Building on our previous findings (unpublished data; preprint submitted), we investigated whether the mode of delivery and the use of lower cell doses, specifically through IN administration of UC-MSCs preconditioned with short severe hypoxia (SSH-MSCs; 0.1% oxygen for 24 hours) or their secretome, can achieve comparable neurofunctional recovery. Thus, two days post-injury, animals received an IN administration of a sub-optimal dose of UC-MSCs (25,000 cells) or the secretome of the cells, and their motor and cognitive function was evaluated at different timepoints of development.

HI-lesioned animals receiving intranasal administration of a suboptimal dose of N-MSCs (HIE+N-MSC group) continued to exhibit significant sensorimotor and cognitive impairments compared to control animals (Figure 6). However, IN administration of SSH-preconditioned UC-MSCs resulted in improvements in sensorimotor function. In the negative geotaxis test, HIE+SSH-MSC animals showed reduced sensorimotor deficits two- (1.4 ± 0.20 seconds, p < 0.0001 vs. HIE; Figure 6A) and five-days (1.36 ± 0.30 seconds, p = 0.002 vs. HIE Figure 6B) following cell administration, demonstrating enhanced reflexes and motor function. The footprint analysis further indicated that these sensorimotor improvements persisted at least until P28, with animals in the HIE+SSH-MSC group exhibiting less overlap (1 ± 0.4 steps, p = 0.0017; Figure 6C) and fewer dragging steps (0 ± 0.4 steps, p = 0.0326; Figure 6D), suggesting sustained functional benefits.

**Figure 6.**
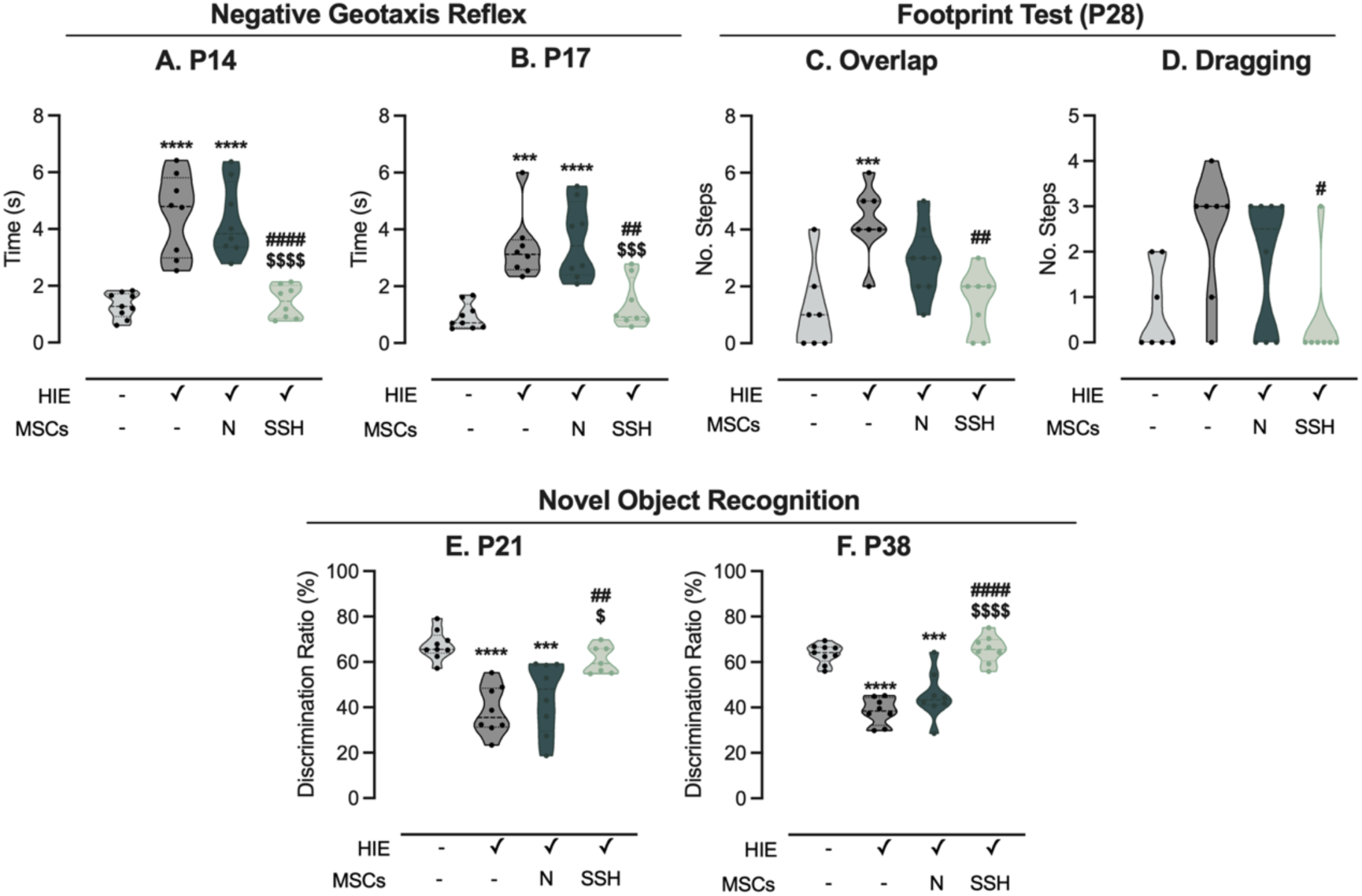
Intranasal administration of hypoxia preconditioned UC-MSCs reduces functional deficits in HI-injured rats. (A, B) Latency (seconds) for pups to rotate 180° and face uphill in the negative geotaxis reflex test at P14 and P17, respectively. (C, D) Footprint analysis assessing dragging and foot overlap at P28, repectively. (E, F) Preference for exploring the novel object (i.e. discrimination ratio) in the novel object recognition test at P21 and P38, respectively. Statistical analysis was performed using one-way ANOVA and Tukey’s multiple comparison tests for the negative geotaxis reflex test and novel object recognition test; and the Kruskal–Wallis test and Dunn’s multiple comparison test for the footprint test. Statistical differences are indicated as: *** p < 0.001 and **** p < 0.0001 vs Control; ^#^ p < 0.05, ^##^ p < 0.01, and ^####^ p < 0.0001 vs HIE; ^$^ p < 0.05, ^$$$^ p < 0.001, and ^$$$$^ p < 0.0001 vs HIE+N-MSC.

Regarding cognitive function, the novel object recognition test assessed recognition memory at P21 (weaning; Figure 6E) and P38 (early adulthood; Figure 6F). At both time points, animals of the HIE+SSH-MSC group displayed a preference for exploring the novel object similar to control animals, indicating an improvement in cognitive outcomes. Moreover, SSH-preconditioned UC-MSCs delivered intranasally were more effective in restoring recognition memory than naïve UC-MSCs, demonstrating the enhanced therapeutic effect of preconditioning (P21: 61 ± 2.3%, p = 0.0238 vs HIE+MN; P38: 66 ± 2.2%, p < 0.0001).

Animals treated with the secretome of SSH-preconditioned UC-MSCs (HIE+SSH-S) showed superior improvements in both sensorimotor and cognitive functions compared to those receiving the secretome of naïve UC-MSCs (HIE+N-S) and untreated HIE animals (Figure 7). In the negative geotaxis test, HIE+SSH-S animals demonstrated faster reorientation, recapitulating the improvement of vestibular reflexes observed in animals treated with the preconditioned cells (Figure 7A, B). Once more, this effect was observable within days of treatment, indicating that this approach was sufficient to induce changes in motor development. The ladder rung walking test revealed sustained motor coordination benefits in HIE+SSH-S animals, with fewer missteps and enhanced gait stability when crossing the apparatus (Figure 7C). These findings contrast with the more frequent missteps and impaired gait pattern seen in HIE and HIE+N-S groups, underscoring the lasting impact of hypoxia-preconditioned factors on balance and gait.

**Figure 7.**
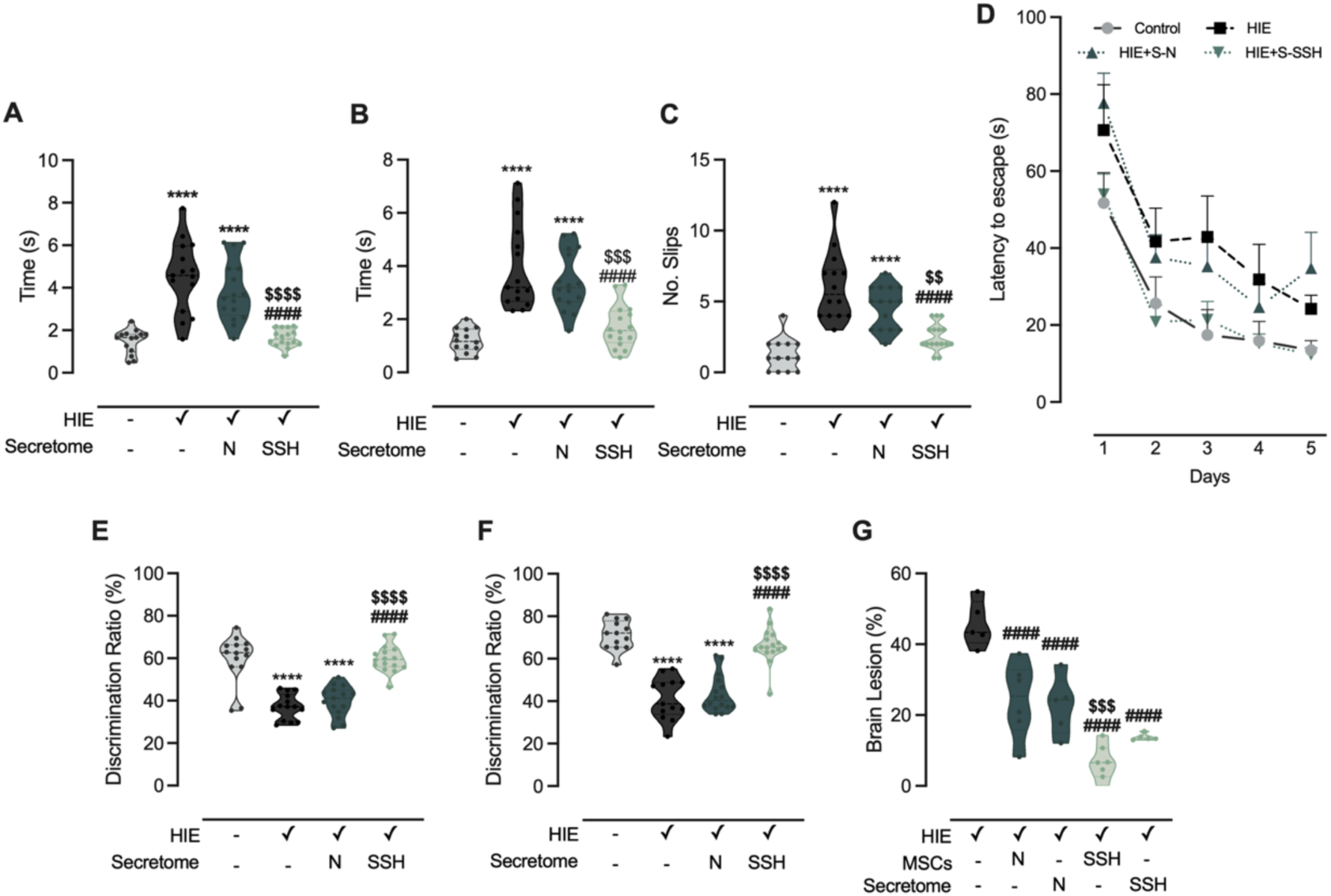
IN-delivery of the secretome from SSH-MSCs promotes neurological recovery in HI-injured rats. (A, B) Latency (seconds) for pups to rotate 180° and face uphill in the negative geotaxis reflex test at P14 and P17, respectively. (C) Number of slips during the ladder rung walking test at P30. (D) Latency to entrance in the escape box (i.e. latency to escape) over five days of testing in the Barnes Maze apparatus. (E, F) Preference for exploring the novel object (i.e. discrimination ratio) in the novel object recognition test at P21 and P38, respectively. (G) Brain lesion extension at P40, measured across sixteen sequential coronal brain sections stained with cresyl violet. Statistical analysis was performed using one-way ANOVA and Tukey’s multiple comparison test for the negative geotaxis reflex test, ladder rung walking test, novel object recognition test, and brain lesion extension determination. Statistical differences are indicated as: *** p < 0.001 and **** p < 0.0001 vs Control; ^####^ p < 0.001 vs HIE; ^$$^ p < 0.01, ^$$$^ p < 0.001, and ^$$$$^ p < 0.0001 vs HIE+N-S (or HIE+N-MSC in G).

Cognitive assessments reflected similar trends. In the Barnes maze, HIE animals treated with the secretome of SSH-preconditioned UC-MSCs consistently located the escape hole quicker than those treated with the naïve secretome or the untreated HIE group, highlighting a marked improvement in spatial learning and memory (Figure 7D). This enhancement in spatial memory retention was sustained throughout the test period, suggesting that the hypoxia-preconditioned secretome elicited robust cognitive recovery. Moreover, in the novel object recognition test, HIE+SSH-S animals displayed a clear preference for the novel object at both weaning (P21) and early adulthood (P38), mirroring control animals and the results obtained with IN administration of SSH-preconditioned UC-MSCs (Figure 7E, D). This finding indicates preserved recognition memory that was enhanced in HIE+SSH-S animals compared to the HIE+N-S group, emphasizing the superior cognitive benefits conferred by the hypoxia-conditioned secretome.

Thirty-days post neonatal HI injury, animals were sacrificed, and sixteen sequential coronal brain sections were stained with cresyl violet to determine the volumes of ipsilesional and contralesional hemispheres and calculate brain lesion extension (Figure 7G). The HIE group presented a brain lesion affecting approximately 45% of the ipsilateral hemisphere, that was significantly reduced in animals treated with IN-delivery of UC-MSCs or their secretome (both naïve and preconditioned with SSH). However, lesioned-animals treated with SSH-preconditioned UC-MSCs (HIE+SSH-MSC group) presented a 74% decrease in brain injury when compared to animals treated with naïve MSCs (p = 0.0002 vs. HIE+ SSH-MSC). Although not significant, the administration of the secretome from the preconditioned cells also showed a tendency to decrease the lesion size when compared to secretome from naïve MSCs (39% decrease vs. HIE+N-S, p = 0.1232).

### 2.4. Long-term glial changes in HIE animals treated with hypoxia-preconditioned UC-MSCs or its secretome delivered intranasally

Building on the previous proteomic findings, which revealed modulation of glial-related proteins following intravenous administration of 50,000 hypoxia-preconditioned UC-MSCs (Chapter 5), we examined whether IN delivery of 25,000 preconditioned UC-MSCs or their secretome could have a long-term impact in glial cells. Using immunohistochemistry, the expression and morphology of GFAP-positive astrocytes and Iba1-positive microglia were assessed in the ipsilesional hemisphere 30 days post-injury.

This analysis revealed distinctive patterns of glial reactivity across experimental groups (Figure 8A). In HI-injured animals, GFAP immunolabeling indicated increased astrocyte reactivity, with a 6.8- and 6.6-fold increase in mean fluorescence intensity (p < 0.0001; Figure 8B) and integrated density (p < 0.0001; Figure 8C) compared to controls. Animals treated with naïve UC-MSCs (HIE+N-MSC) and SSH-preconditioned UC-MSCs (HIE+SSH-MSCs) had reduced GFAP mean intensity (p < 0.0001 vs. HIE) and integrated density (p < 0.0001 vs. HIE), suggesting attenuation of astrocyte reactivity following stem cell administration. Treatment with the secretome yielded similar results. However, hypoxia-preconditioned UC-MSCs or their secretome further lowered GFAP mean intensity (p < 0.0001 HIE+N-MSC vs. HIE+SSH-MSC; p = 0.0006 HIE+N-S vs. HIE+SSH-S; Figure 8B) and integrated density (p < 0.0001 HIE+N-MSC vs. HIE+SSH-MSC; p = 0.0043 HIE+N-S vs. HIE+SSH-S; Figure 8B) compared to administration of naïve UC-MSCs. These changes mirror previous proteomic results, where IV administration of SSH-preconditioned UC-MSCs normalized GFAP levels more effectively than naïve UC-MSCs administration (Figure 8).

**Figure 8.**
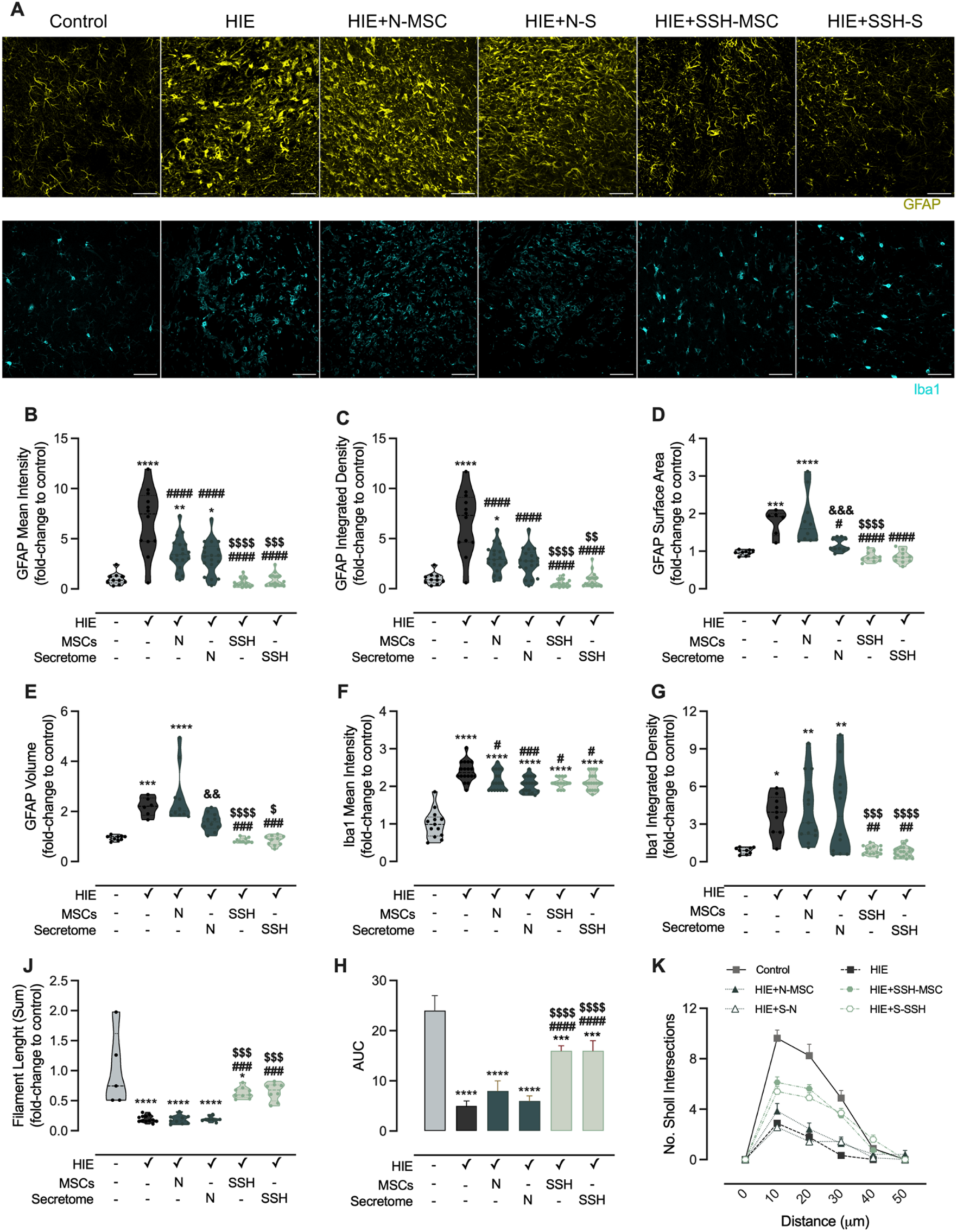
Administration of SSH-preconditioned UC-MSCs or its secretome modulated long term glial response in HIE rats. (A) Representative brain sections from P40 Wistar rats labelled for GFAP or Iba1 acquired with a 40× objective using the z-stack function (scale bar = 50 µm). (B, C) GFAP mean intensity and integrated density expressed as fold-change to control, respectively. (D, E) GFAP surface area and volume with 3D surface rendering expressed as fold-change to control, respectively. (F, G) Iba1 mean intensity and integrated density expressed as fold-change to control, respectively. (J, H) Evaluation of microglia branching by the sum of filament length per cell as fold-change to control and area under the curve for each experimental group, respectively, in (K) Sholl analysis. Statistical differences between groups are represented in the figures with * p < 0.05, ** p < 0.01, *** p < 0.001, **** p < 0.0001 vs. control; ^#^ p < 0.05, ^##^ p < 0.01, ^###^ p < 0.001, ^####^ p < 0.0001 vs. HIE; ^$^ p < 0.05, ^$$$^ p < 0.001, ^$$$$^ p < 0.0001 HIE+N-MSC vs. HIE+SSH-MSC or HIE+N-S vs. HIE+SSH-S; ^&&^ p < 0.01, ^&&&^ p < 0.001 HIE+N-MSC vs. HIE+N-S. Abbreviations: AUC: area under the curve; No.: number.

The morphological analysis of GFAP-positive cells using 3D surface rendering provided additional insights into astrocyte changes. HI injury triggered a 1.8-fold increase in GFAP-positive cell surface area (p = 0.0003 vs. control; Figure 8D) and a 2.2-fold increase in cell volume (p = 0.0005 vs. control; Figure 9E), indicating hypertrophic changes characteristic of reactive astrocytes. Treatment with SSH-preconditioned UC-MSCs or their secretome normalized these parameters to control levels (Figure 8D, E), while naïve UC-MSCs showed limited impact on astrocyte hypertrophy. The secretome from naïve UC-MSCs (HIE+N-S group) had an intermediate effect, reducing GFAP-positive surface area (p = 0.0007 vs. HIE+N-MSC) and volume (p = 0.0015 vs. HIE+N-MSC) to a greater degree than the administration of naïve UC-MSCs.

**Figure 9.**
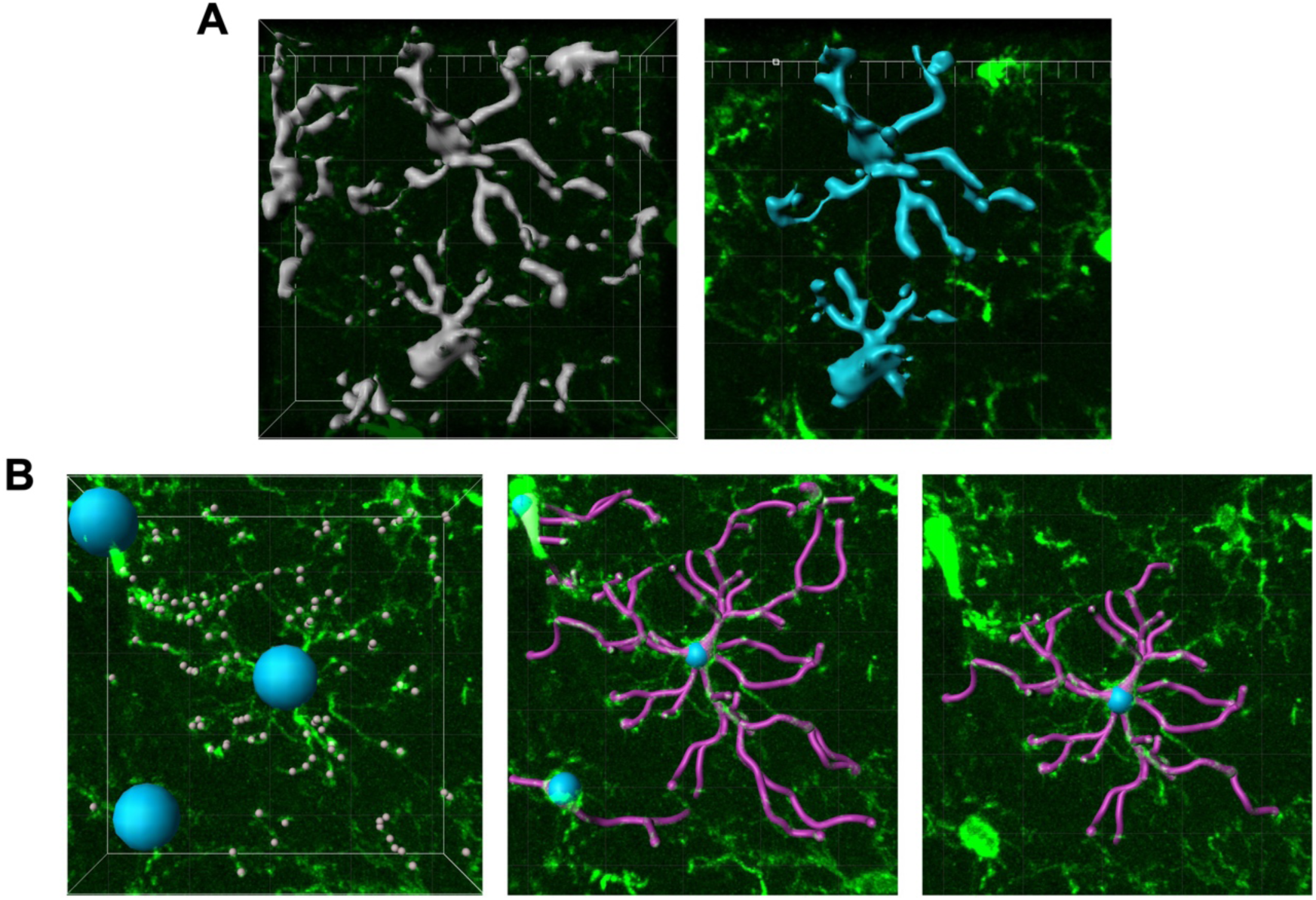
Morphological analysis of microglia using Imaris. The surface of the microglia in the ROI was rendered using the “Surface” function. Automatic settings were used to define the microglial cells. The same ROI was used for the segmentation of each microglial process using the “Filament” function. The automatic protocol for segmentation of the surfaces (A) and filaments (B) that constitute true microglial cells was manually adjusted to create a protocol that suits all experimental conditions. After creating the surfaces and filaments, the disconnected segments and incomplete microglia were manually deleted.

Microglial activation, assessed through Iba1 immunolabeling, further highlighted treatment-dependent effects. Iba1 expression significantly increased in HIE animals, compared to controls (2.4-fold change in mean intensity, p < 0.0001 vs. control; 3.8-fold change in integrated density, p = 0.0193 vs. control; Figure 8F, G). IN administration of SSH-preconditioned or naïve UC-MSCs, or their secretomes, reduced Iba1 mean intensity compared to untreated-HIE animals (Figure 8F). However, the two groups treated with an approach involving hypoxic preconditioning, showed decreased Iba1 integrated density levels versus HIE (p = 0.0069 and p = 0.0018 vs. HIE, respectively; Figure 8G), and the two non-preconditioned therapy groups (p = 0.0002 HIE+N-MSC vs. HIE+SSH-MSC; p < 0.0001 HIE+N-S vs. HIE+SSH-S; Figure 8G).

Detailed morphological analysis of microglial cells using filament tracing and Sholl analysis further supported the potential of administering UC-MSCs preconditioned with SSH-preconditioned to modulate glial cells upon HI injury. HIE group presented reduced microglial branching complexity (0.2-fold change to control; p < 0.0001: Figure 8J), indicative of a reactive phenotype with simplified morphology. Treatment with SSH-preconditioned UC-MSCs or their secretome restored microglial ramification toward control levels (p = 0.0002 and p = 0.0002 vs. HIE, respectively), suggesting a shift to a more regulatory, neuroreparative state. This interpretation is reinforced by Sholl analysis, where the area under the curve for groups treated with hypoxia-preconditioned therapy is closer to control animals (Figure 8H, K), and higher to that of HIE and groups treated with non-preconditioned therapies.

## 3. Discussion

This study evaluates three key strategies to potentiate the use of UC-MSC in neonatal HIE: the comparative effects of IV versus IN administration, the role of hypoxic preconditioning, and the therapeutic potential of the secretome derived from preconditioned UC-MSCs. IN delivery demonstrated distinct advantages when compared with IV administration, with IN administration requiring fewer cells to achieve comparable functional recovery and glial modulation. On the other hand, hypoxic preconditioning further enhanced the efficacy of UC-MSCs, amplifying their ability to attenuate neurological impairments while decreasing astrocyte and microglial reactivity. Also, the secretome from these preconditioned cells replicated the therapeutic effects, underscoring its potential as an acellular alternative that retains the full spectrum of bioactive molecules for modulating inflammatory and repair pathways. Together, these findings highlight the versatility and optimization potential of UC-MSC-based therapies for neonatal HIE.

First, we aimed to evaluate the influence of the UC-MSCs administration route on the extension of brain damage and neurological outcomes after a HI insult in postnatal Wistar rats modeling neonatal HIE. For that purpose, two days after induction of the HI brain lesion, the animals were treated with a low dose of UC-MSCs (50,000) delivered intranasally or intravenously. The main hypothesis of this study was that IN administration would result in superior outcomes compared to IV administration, when using a low UC-MSCs dose, due to its potential for direct delivery to the brain and reduced systemic dispersion and trapping.

The IN-administration route has been shown to offer several advantages, including increased cell targeting to the lesion site, non-invasiveness, efficiency, and minimal risk of infection (9). This route has gained increasing interest for delivering therapeutic agents to the brain due to the anatomical connection, which facilitates the transportation of substances directly to the central nervous system, bypassing the blood-brain barrier (10). A phase-I clinical study reported that IN-delivered bone marrow MSC in neonates who suffered a perinatal arterial ischemic stroke was safe and feasible (11). The present study found that IN delivery of MSCs resulted in a smaller brain lesion volume compared to both the untreated HI group and the intravenously treated group. Additionally, the improvements in gait coordination, balance, and cognitive function were more pronounced in the IN-administration group compared to the IV-administration group. This greater efficacy correlates with a higher capacity to reverse the glial alterations caused by hypoxic-ischemic injury, namely in reducing astrocyte and microglia reactivity, and increasing myelination, proposing glia modulation of glial as a potential mechanism underlying the brain repair and functional recovery in HIE injured animals treated with UC-MSCs. This may also indicate that the therapeutic potential of intravenously delivered UC-MSCs reported in previous studies was due to the higher dose used (0.1 to 6 × 10^6^ cells) (12–16). Indeed, our data indicates that the administration of 0.5 × 10^6^ UC-MSCs by this route was able to promote functional recovery of this model of HIE (Supplementary Figure 1). Thus, the literature and the findings present in this study support the idea that IN-delivery of UC-MSCs requires fewer cells to effectively promote neurological recovery after neonatal HI injury, when compared with the IV route. The impact of the delivery route on the effective dose may be related with the low migration rate of MSCs to the lesion site when administered intravenously. When compared with IN administration, the compromised homing and survival of MSCs in IV delivery justifies the need of higher doses to achieve therapeutic efficacy (17).

While most of the studies evaluating the therapeutic potential of IN-administration of MSCs in preclinical models of HIE report that this strategy is effective in reducing functional impairments and brain lesions, these studies often use considerably higher doses of MSCs when compared to the present study (18, 19). For instance, McDonald et al (2019) reported that the IN administration of 0.2×10^6^ UC-MSCs following HI was neuroprotective and reduced brain inflammation (19). The results of the present study are in concordance with this conclusion, even with a 75% reduction of administered cells.

Additionally, IN administration of UC-MSCs effectively modulated glial reactivity, which plays a critical role in neuroinflammation and tissue repair. Most likely, MSCs can attenuate neuroinflammation and promote tissue healing due to a prompting a shift from a pro-inflammatory to an anti-inflammatory phenotype in glial cells. Moreover, findings from previous studies suggest that neuroinflammation, namely activated microglia induced by hypoxia-ischemia, inhibits oligodendrocyte maturation (20, 21). By sustaining increased levels of pro-inflammatory mediators after a hypoxic-ischemic insult, including TNF-α, IL-1β, IL-6, and complement pathways, the reactive microglia promote white matter damage (22). The results obtained in the present study showed that decreased microglial reactivity induced by IN administration of UC-MSCs was correlated with increased myelinization of the corpus callosum and increased MBP-structural complexity. This data supports previous findings achieved with doses of IN-administered MSCs up to 20-times higher (23). By understanding how MSCs modulate neuroinflammatory processes and interact with key signaling pathways involved in brain injury, treatment strategies can be tailored aiming at mitigating neonatal brain damage, promoting repair, and ultimately improving long-term neurodevelopmental outcomes. The administration of hypoxia-preconditioned UC-MSCs and their secretome exerted similar effects on both astrocyte and microglial responses in HI-injured animals. GFAP immunolabeling revealed that hypoxia-preconditioned treatments reduced astrocyte reactivity more effectively than naïve UC-MSCs, as evidenced by lower mean fluorescence intensity and integrated density, alongside normalization of hypertrophic changes in astrocytes to control levels. The analysis of microglia reactivity also suggested treatment-dependent effects. While all treatments reduced Iba1 expression per cell, only hypoxia-preconditioned therapies lowered the overall number of microglia cells in the perilesional area, as indicated by reductions in Iba1 integrated density. Morphological assessments further emphasized the efficacy of hypoxia-preconditioned UC-MSCs in restoring microglial complexity and ramification, shifting microglial phenotype toward a regulatory, neuroreparative state. Research over the past few decades has highlighted critical functions of microglia in brain development, including synaptic pruning, maintenance of brain structure, clearance of cell debris, axon guidance, and the regulation of neurogenesis and oligodendrogenesis [reviewed in Chapter 1, (24)]. Following HI brain injury in the developing brain, microglia undergo rapid phenotypical changes, adopting an amoeboid morphology associated with microglial activation and expression of OX-6, OX-18 and CD68 markers (25). Studies using animal models of neonatal HIE indicate that microglial activation varies by brain region, with the microglia in the hippocampus being first activated after injury, followed by activation of cortical and striatal microglia (26). Over time, microglia appear to express both pro- and anti-inflammatory markers, challenging the traditional M1/M2 paradigm (27, 28). Similarly, our study and prior preclinical research show that microglia maintain persistent phenotypic changes during the tertiary phase of injury (7, 23, 26–29). Brégère et. al. (2021) recently reviewed studies linking stem cell therapy with neurological improvements and modulation of microglial activation (25). This analysis revealed variability in MSCs’ effect on microglia in HIE models, though most studies reported that MSCs reduced microglial activation (primarily measured using Iba1 and CD68 markers) correlated with behavioral improvements and limited neuroprotection (25). Identified discrepancies could stem from differences in stem cell potency or from suboptimal dosing that failed to induce a consistent effect. Our study demonstrated a reliable, beneficial modulation of the long-term microglial response in lesioned animals treated with low doses of hypoxia-preconditioned UC-MSCs delivered by two different routes of administration – IV and IN. We combined fluorescence intensity and integrated density measurements of Iba1 labeling with morphological assessments, including Sholl analysis, to capture a comprehensive view of microglia phenotype. This multifaceted approach provides a deeper insight into microglial modulation after stem cell therapy in HIE models. Our findings suggest that both naïve and SSH-MSCs reduced mean Iba1 intensity; however, only SSH-preconditioned treatments lowered Iba1 integrated density. This particular outcome may help to explain conflicting results in previous studies, pointing out that only partial aspects of the microglial response were previously captured. Our comprehensive analysis shows that assessing multiple parameters can provide a more complete picture of treatment effects on microglia response after injury.

Emerging evidence positions the MSC secretome as a promising acellular alternative to live cell therapy. In this study, IN administration of the secretome from SSH-MSCs replicated the functional and glial outcomes of cell-based treatments using the same preconditioning strategy. These observations suggest that hypoxic preconditioning primes UC-MSCs to release factors that more effectively modulate inflammatory and repair pathways. To our knowledge, few studies have explored the therapeutic potential of the secretome derived from stem cells in neonatal HIE models, and none have evaluated the impact of IN administration of the secretome from SSH-preconditioned UC-MSCs. Wei et. al. first reported that secretome from adipose-derived stem cells reduced brain lesion size and improved cognitive function in a rodent HIE model (30). Recently, Huang et. al., found that intranasal delivery of secretome from human pluripotent stem cell-derived ectomesenchymal stromal cells reduced brain lesion size while decreasing GFAP and Iba1 mean intensity (31). Additionally, administration of MSC-derived extracellular vesicles, an isolated fraction of secretome, was reported to induce significant functional improvement in animal models of HIE (31–36), while reducing Iba1-positive area fractions and Iba1-positive cells in the cortex and corpus callosum (32, 33, 36). Nonetheless, using complete secretome - as in our study – can offer key advantages over isolated extracellular vesicles. The complete secretome retains the full spectrum of bioactive molecules, including soluble factors, metabolites, and extracellular vesicles, thus preserving their natural stoichiometry and interactions (37, 38). This complex biological profile more closely mimics the physiological paracrine signaling environment and can enable synergistic effects that boost therapeutic potential. Also, complete secretome preparation involves fewer processing steps, reducing potential alterations from manipulation and providing higher yields of therapeutic material. Isolated extracellular vesicles, though effective, may lack beneficial non-vesicular factors and disrupt the natural paracrine signaling balance. Recent studies comparing isolated extracellular vesicles to complete secretome have shown differential effects that could impact immunomodulatory potential, favoring complete secretome as a more advantageous option (39, 40).

Nonetheless, it is still necessary to perform an exhaustive characterization of the mechanisms underlying the neurological recovery observed with the administration of UC-MSCs (preconditioned or not) and its targets to advance stem cell-based therapies to clinical applications.

## 4. Conclusion

Our study demonstrated greater efficacy in adopting IN administration of UC-MSCs to reduce brain lesions and deficits associated with neonatal HI brain injury when compared to IV delivery, while also improving motor coordination, recognition memory, glial cell response, and myelination in the corpus callosum. Furthermore, combination of intranasal administration and UC-MSCs’ preconditioning reduced significantly the dose of cells required to achieve neurological recovery and modulation of glial responses, a hallmark of the neonatal HIE chronic phase, making this approach more accessible for clinical use. Notably, the secretome from preconditioned cells replicated the reparative effects of cellular therapy, highlighting the potential of acellular approaches in treating neonatal HIE. These findings underscore the importance of optimizing MSC therapy for neonatal neurorepair and promotion of long-term cognitive and motor outcomes in affected individuals.

## 5. Methods

### 5.1. Ethical Approval

Approval for animal studies was obtained from the Ethical Committee of the University of Beira Interior and authorized by the Portuguese General Directorate for Food and Veterinary (0421/000/000/2019). The research adhered to the European Directive (2010/63/EU) governing the protection of laboratory animals used for scientific purposes. Additionally, ethical approval for using human mesenchymal stem cells was granted by the Ethical Committee of the Faculty of Medicine of the University of Coimbra (Approval Number: 075-CE-2019). The use of animals in this project is justified by the importance of assessing the impact of the proposed strategies on functional outcomes, such as cognitive and motor capabilities, for their translation to clinical applications. The animal research conducted in this study is reported following the Animal Research: Reporting of In Vivo Experiments (ARRIVE) 2.0 guidelines (41).

### 5.2. Animals

Animal studies were carried out in the animal facility of the Faculty of Health Sciences, University of Beira Interior (Covilhã, Portugal). Animals enrolled in this study were maintained in an alternating 12-hour light/dark cycle, maintained with their dam until weaning at P21, and handled daily after the induction of the neonatal HI injury to monitor welfare. Experimental design, including sample size calculation, was done using Experimental Design Assistant (https://eda.nc3rs.org.uk). The effect size and variability were determined by previous experiments from the group. The individual animal served as the experimental unit and was randomly and independently assigned to one of the experimental groups with sex as a blocking factor. The researchers were not blinded for the behavioral tests or statistical analysis.

### 5.3. Induction of neonatal HI brain injury

Male and female Wistar rat pups at P10 (weighting 15-20 g) were subjected to an HI brain lesion following the adapted Rice-Vannucci model for HIE. First, the animals were anesthetized using isoflurane (5% for induction, 1.5-2% for maintenance; Isoflo, Zoetis). Them, the left common carotid artery was exposed and ligated using 6−0 silk suture (F.S.T).

For the hypoxia, the animals were placed in an airtight chamber at 37°C filled with a mixture of 8% oxygen and 92% nitrogen (Air Liquide) for 90 minutes. Control animals underwent a similar procedure; however, their left common carotid artery was not ligated and were exposed to room air in a heated chamber at 37°C.

### 5.4. UC-MSCs culture

UC-MSCs were isolated from the Wharton’s Jelly of cryopreserved fragments of human UC, as previously described (42). When colonies were observed, cells were washed with PBS, detached by the addition of 0.05% Trypsin-EDTA (Gibco™) on a humidified incubator at 37°C. Then, UC-MSCs were homogenized and centrifuged at 290×g, at room temperature. The cell pellet was resuspended and UC-MSCs were replated in Minimum essential Medium-α (Gibco™) supplemented with 5% (v/v) fibrinogen depleted human platelet lysate (HPL) (UltraGRO™, Helios) and Antibiotic-Antimycotic (Gibco™), until sub confluence was achieved. This procedure was repeated until passage four and UC-MSCs were cryopreserved in HPL with 10% dimethyl sulfoxide.

### 5.5. Hypoxic preconditioning of UC-MSCs and secretome preparation

UC-MSCs were plated in Minimum Essential Medium-α (Gibco™) supplemented with 5% (v/v) fibrinogen depleted HPL (UltraGRO™, Helios) and Antibiotic-Antimycotic (Gibco™), until sub confluence was achieved. This procedure was repeated to expand the number of cells in culture. For hypoxic preconditioning, UC-MSCs were kept at standard culture conditions until passage three. Then, 24 hours before the stimuli, cells were seeded at 10,000/cm^2^ (i.e. passage four). Before the stimuli, the culture media was washed with PBS and replaced with Minimum Essential Medium-α (Gibco™) without supplementation, and UC-MSCs were placed for 24 hours on a humidified incubator (Binder), at 37°C, with 5% CO2 (N-MSCs) – or in an InvivO₂® 400 humidified incubator (Baker Ruskinn), set at 0.1% O_2_/5% CO_2_ (SSH-MSCs) or 5% O_2_/5% CO_2_ (MH-MSCs). After the stimuli, the culture media was collected and centrifuged at 290×g, at room temperature, to remove cell debris. Then, the supernatant was transferred to a low molecular weight cut-off (5kDa Vivaspin® 20, Sartorius) and centrifuged at 3,000×g, at 4°C, until the desired concentration was achieved (about 50x). UC-MSCs were cryopreserved in HPL with 10% dimethyl sulfoxide and stored in liquid nitrogen until administration.

### 5.6. UC-MSCs and secretome administration

MSCs were delivered by IV or IN route two days after the induction of HI brain lesion (i.e. P12). On the day of administration, the previously prepared MSCs were thawed and centrifuged at 290×g for 5 minutes. After removing the supernatant, MSCs were resuspended in PBS and viable cellular density was determined using the Trypan Blue exclusion method.

For IV administration, the animals were anesthetized with isoflurane and 50,000 viable UC-MSCs diluted in 200 µl of PBS were administered, per animal, in the tail vein using a 29-gauge insulin syringe (Terumo).

For IN administration, 25,000 or 50,000 viable UC-MSCs/iMSCs diluted in 24 µl of PBS were administered per animal. The administration was performed in two moments with an interval of two hours. In each moment of administration, the animals were immobilized in a supine position and two microliters of the MSCs suspension was administered six times to each nostril using a micropipette, with an interval of five minutes between each administration and alternating amongst nostrils.

The secretome was administered in a series of six periods of administration per animal, separated by two-hour intervals. Each dose consisted of two microliters of secretome, administered into alternating nostrils using a micropipette. The five-minute wait between administrations was crucial for optimal absorption, and the animal was held in a supine position for ten seconds afterwards. Control animals received the same volume of PBS using the same administration technique to maintain consistency across the experiment. The amount of secretome administered corresponded to 0.5 x 10^6^ UC-MSCs.

### 5.7. Behavioral analysis

#### 5.7.1. Negative Geotaxis Reflex

The negative geotaxis reflex test was used to assess rat’s motor coordination early in development, at P14 and P17. For this test, rat pups were placed downhill on a 45° slanted slope, and the time required for the pups to face uphill was recorded. No animals were excluded from the analysis.

#### 5.7.2. Novel Object Recognition Test

The recognition memory of the animals was assessed using the novel object recognition test at two developmental stages: infancy (P21) and early adulthood (P38) (43). Prior to the test day, animals underwent a habituation phase in the testing arena, lasting 10 minutes. On the test day, animals were initially presented with two identical objects during a 10-minute familiarization phase. After a 30-minute interval, the animals were exposed to one familiar object and one novel object for 10 minutes during the test phase. Exploration times for both the familiar and novel objects were recorded during the initial five minutes of the test phase. Subsequently, a discrimination ratio was computed using the formula: (time spent exploring the novel object) / (total exploration time) × 100. A similar procedure was followed for the assessment at P38, except that different sets of familiar and novel objects were used.

#### 5.7.3. Footprint test

As previously described, the footprint test served as the primary method for identifying locomotor deficits and gait abnormalities (44, 45). The assessment was conducted at P28, corresponding to a developmental stage akin to human childhood (43). Rats’ fore and hind paws were coated with non-toxic paint, following which they were prompted to cross a 100 cm path lined with paper, leading towards a black box. Evaluation of the footprint pattern entailed analyzing ten consecutive steps, during which instances of foot-dragging or overlapping footprints of the contralesional paws were counted. Animals showing a lack of motivation to cross the corridor were excluded from subsequent analysis.

#### 5.7.4. Ladder Rung Walking Test

The assessment of animals’ motor function, particularly coordination, was conducted using the ladder rung walking test at P30, a developmental stage corresponding to human childhood (43). The apparatus for this test consisted of a 100 cm long corridor delineated by two transparent acrylic side walls, between which metal rods (0.3 cm in diameter) were positioned 1 cm apart. Rats crossed the apparatus four consecutive times under video recording. Subsequently, each video was reviewed in slow motion to count the number of foot slips occurring between the rods. Animals showing a lack of motivation to cross the ladder were excluded from subsequent analysis.

#### 5.7.5. Barnes Maze Test

The Barnes maze test was applied to evaluate spatial memory and learning. The apparatus consisted of a round table with 23 holes, with one of the holes leading to an escape box. This apparatus was set in a well-lit room with four visual cues placed in a fixed position throughout all testing days. The testing began at P31, with the habituation phase followed by the first day of the acquisition phase. During habituation, the rat was placed inside the escape box for one minute and afterwards was allowed to explore the apparatus freely; if the rat failed to find the escape hole within five minutes, it was guided to it. After entering the escape box, the animal was maintained there for 30 seconds. The acquisition phase consisted of two trials per day, with an inter-trial interval of 20 minutes, for five consecutive days. For each trial, the rat was placed in the center of the apparatus and allowed to explore for a maximum period of three minutes. The test was recorded with a video camera using the ANY-Maze software (Stoelting).

### 5.8. Tissue Collection and Preparation

At postnatal day 40, the animals were euthanized with an overdose of anesthetics (200 mg/kg Ketamine and 10 mg/kg Xylazine), followed by perfusion with PBS and 4% paraformaldehyde. Following perfusion, the brains were excised and immersed in 4% paraformaldehyde for 16 hours at 4 °C. Subsequently, for cryopreservation purposes, the brains were transferred to a 30% sucrose solution until they sank. Once cryopreserved, the brain tissue was rapidly frozen in liquid nitrogen and stored at −80 °C until further processing.

Sectioning of the frozen brains was performed using a cryostat (Leica CM3050) set to a thickness of 40 µm, with sections collected at 240 µm intervals.

### 5.9. Cresyl Violet Staining

Brain lesion extension was assessed in sixteen sequential brain sections for each animal. Frozen brain sections were mounted in poly-lysin coated glass slides (VWR) and stained with 0.05% Cresyl Violet Acetate (Merck) using the Sakura TissueTek Automated DRS 2000 automated slide stainer. Images were acquired with the Axio Imager A1 Microscope (Zeiss) with a 5× objective (EC Plan-Neofluar 4×/0.16 M27), and the volume of the left (ipsilesional) and right (contralesional) hemispheres were determined with the Cavalieri’s Principle Probe of the StereoInvestigator software (MBF Bioscience). Brain lesion extension was calculated as (V_contralesional_ − V_ipsilesional_)/V_contralesional_ × 100.

### 5.10. Immunohistochemistry

Brain sections underwent permeabilization with PBS-1% Triton-X100 for 30 minutes, followed by blocking for 2 hours using 10% fetal bovine serum (Biochrom) and 0.1% Triton-X100. Subsequently, sections were incubated for 72 hours at 4°C with rabbit anti-GFAP antibody (DAKO Z0334, 1:200) or rabbit anti-ionized calcium binding adaptor molecule 1 (Iba1) antibody (WAKO 019-19741, 1:2000). For myelinization analysis, sections were incubated for 40 hours at 4°C with a mouse anti-myelin basic protein (MBP) antibody (Biolegend 836504, 1:200).

Following primary antibody incubation, sections were incubated for 2 hours at room temperature with the corresponding secondary antibodies using a 1:1000 dilution: anti-rabbit A488 (Molecular Probes A11008) or anti-mouse A488 (Invitrogen A11001). Nuclei were counterstained with Hoechst 33342 (Invitrogen H1399, 1:1000).

### 5.11. Image Acquisition and Processing

For GFAP and Iba1 labeling, images of the peri-infarct area were acquired using the LSM 710 AxioObserver Microscope (Zeiss) with a 20× objective (Plan-Apochromat 20×/0.8 M27) or 40× objective (EC Plan-Neofluar 40×/1.30 Oil DIC M27) in four adjacent tiles for the quantification of mean fluorescence intensity and integrated density. GFAP signal was captured with the following settings, emission/excitation wavelength: 561/488 and Pinhole: 1 AU. Iba1 signal was captured using the following settings, emission/excitation wavelength: 561/488 and Pinhole: 1 AU. For glia morphological analysis, confocal 3D images were acquired using the z-stack function. Images were acquired at 0.61 µm interval in LSM 710 AxioObserver Microscope (Zeiss) with a 40× objective (EC Plan-Neofluar 40×/1.30 Oil DIC M27).

For MBP labeling, images of the corpus callosum (approximate coordinates: − 2.8 < y < −2.5 mm from bregma(46)) were acquired with the 20× objective in six adjacent tiles. MBP signal was captured using the following settings, emission/excitation wavelength: 561/488; and Pinhole: 1 AU.

#### 5.11.1. Quantification of fluorescence intensity

Mean fluorescence intensity and the integrated density of GFAP and Iba1 labeling were quantified with ImageJ software in four non-overlapping fields of view of the peri-infarct area (or equivalent region in the control group) per section in three to four sections per animal. MBP fluorescence intensity was measured in the micrographs with all acquired tiles for each section.

#### 5.11.2. Characterization of oligodendrocytes and myelin staining

The MBP-positive area was determined in ImageJ by defining a threshold that identifies only MBP labeling and measuring the proportion of the acquired field that was labeled (% Area). The coherency of MBP-positive fibers was determined similarly to Tilborg *et al.* (2017) (47) using ImageJ plugin OrientationJ (8) in a selected region of interest of the corpus callosum. This parameter is an inverse measure of complexity of white matter microstructure; thus, a more complex organization will have decreased coherency between myelinated structures.

#### 5.11.3. Morphological analysis of glial cells

Morphological analysis was performed on 3D images using Imaris 7.6.4 software (Bitplane, Zurich, Switzerland). The analysis was adapted from a published protocol (48) and is exemplified in Figure 9.

### 5.12. Statistical Analysis

Statistical analysis was performed using GraphPad Prism 10 (GraphPad Software Inc., San Diego, CA). Outliers from each data set were identified with the Grubbs’ analysis (α=0.05) and excluded from the analysis. Statistical analysis was performed with the Kruskal Wallis test coupled with Dunn’s multiple comparison correction test or One-way ANOVA coupled with Tukey’s multiple comparison test, according to the type of data. The differences were considered significant when the p-value was inferior to 0.05. Details of the statistical analysis can be found in the figure legends.

## Notes

### Competing Interest Statement

The authors have declared no competing interest.

