## Supplementary material for "Nose-to-Brain Healing: Hypoxia-Preconditioned Mesenchymal Stem Cells Prompt Recovery in Hypoxic-Ischemic Encephalopathy Rats": Table S1

**Table S1.** Statistical analysis performed for each experiment included in this study.

| Figure Number | Experimental Condition | N | Mean | SEM | One-Way ANOVA Test Value and Degrees of Freedom | One-Way ANOVA p-value | Tukey's Multiple Comparisons Post-Hoc p-values | | |
|---|---|---|---|---|---|---|---|---|---|
| | | | | | | | vs Control | vs HIE | vs HIE+MSC_IV |
| 1 | Control | 6 | 0.0030 | 0.0017 | F (3, 20) = 33.32 | < 0.0001 | | | |
| | HIE | 6 | 0.4374 | 0.0147 | | | < 0.0001 | | |
| | HIE+MSC_IV | 6 | 0.4595 | 0.0577 | | | < 0.0001 | 0.9771 | |
| | HIE+MSC_IN | 6 | 0.1476 | 0.0495 | | | 0.0689 | 0.0002 | < 0.0001 |
| 2A | Control | 12 | 1.438 | 0.176 | F (3, 43) = 11.71 | < 0.0001 | | | |
| | HIE | 12 | 4.143 | 0.436 | | | < 0.0001 | | |
| | HIE+MSC_IV | 12 | 3.768 | 0.175 | | | 0.0001 | 0.8727 | |
| | HIE+MSC_IN | 11 | 3.154 | 0.522 | | | 0.0079 | 0.2203 | 0.6206 |
| 2B | Control | 12 | 1.203 | 0.140 | F (3, 43) = 11.82 | < 0.0001 | | | |
| | HIE | 12 | 3.752 | 0.499 | | | < 0.0001 | | |
| | HIE+MSC_IV | 12 | 3.847 | 0.369 | | | < 0.0001 | 0.9978 | |
| | HIE+MSC_IN | 11 | 2.295 | 0.383 | | | 0.1845 | 0.0422 | 0.0272 |
| 2C | Control | 11 | 1.2 | 0.4 | F (3, 40) = 11.23 | < 0.0001 | | | |
| | HIE | 11 | 5.9 | 0.8 | | | < 0.0001 | | |
| | HIE+MSC_IV | 11 | 4.5 | 0.4 | | | 0.0019 | 0.3178 | |
| | HIE+MSC_IN | 11 | 3.6 | 0.7 | | | 0.0355 | 0.0355 | 0.6996 |
| 2D | Control | 12 | 0.5 | 0.2 | F (3, 44) = 6.369 | 0.0011 | | | |
| | HIE | 11 | 2.7 | 0.6 | | | 0.004 | | |
| | HIE+MSC_IV | 14 | 2.6 | 0.4 | | | 0.0046 | 0.9935 | |
| | HIE+MSC_IN | 11 | 2.6 | 0.5 | | | 0.0061 | 0.9989 | 0.9995 |
| 2E | Control | 12 | 0.5 | 0.2 | F (3, 44) = 6.312 | 0.0012 | | | |
| | HIE | 11 | 2.0 | 0.4 | | | 0.0121 | | |
| | HIE+MSC_IV | 14 | 1.9 | 0.4 | | | 0.0111 | 0.9985 | |
| | HIE+MSC_IN | 11 | 0.6 | 0.2 | | | 0.9911 | 0.0305 | 0.0299 |
| 2F | Control | 11 | 70.55 | 2.21 | F (3, 40) = 24.86 | < 0.0001 | | | |
| | HIE | 11 | 42.23 | 3.13 | | | < 0.0001 | | |
| | HIE+MSC_IV | 11 | 42.65 | 3.61 | | | < 0.0001 | 0.9996 | |
| | HIE+MSC_IN | 11 | 63.0 | 2.37 | | | 0.2656 | < 0.0001 | < 0.0001 |
| 2G | Control | 12 | 61.61 | 2.82 | F (3, 43) = 17.97 | < 0.0001 | | | |
| | HIE | 12 | 37.80 | 1.71 | | | < 0.0001 | | |
| | HIE+MSC_IV | 12 | 44.83 | 3.16 | | | 0.0001 | 0.2129 | |
| | HIE+MSC_IN | 11 | 55.77 | 2.14 | | | 0.3865 | < 0.0001 | 0.0218 |
| 3B | Control | 24 | 695129 | 126953 | F (3, 91) = 34.03 | < 0.0001 | | | |
| | HIE | 24 | 5750617 | 663056 | | | < 0.0001 | | |
| | HIE+MSC_IV | 24 | 3598003 | 471583 | | | < 0.0001 | 0.0025 | |
| | HIE+MSC_IN | 23 | 739277 | 115464 | | | 0.9999 | < 0.0001 | < 0.0001 |
| 3C | Control | 24 | 10.08 | 0.37 | F (3, 91) = 26.02 | < 0.0001 | | | |
| | HIE | 24 | 18.96 | 1.22 | | | < 0.0001 | | |
| | HIE+MSC_IV | 24 | 15.33 | 0.96 | | | 0.0001 | 0.0126 | |
| | HIE+MSC_IN | 23 | 10.61 | 0.37 | | | 0.9698 | < 0.0001 | 0.0007 |
| 3D | Control | 24 | 1098355 | 129600 | F (3, 91) = 17.07 | < 0.0001 | | | |
| | HIE | 24 | 1098355 | 655484 | | | < 0.0001 | | |
| | HIE+MSC_IV | 23 | 3204485 | 476215 | | | 0.0028 | 0.1482 | |
| | HIE+MSC_IN | 24 | 981029 | 76542 | | | 0.9970 | < 0.0001 | 0.0014 |
| 3E | Control | 24 | 19.67 | 0.59 | F (3, 92) = 15.97 | < 0.0001 | | | |
| | HIE | 24 | 24.75 | 1.06 | | | 0.0006 | | |
| | HIE+MSC_IV | 24 | 28.25 | 1.13 | | | < 0.0001 | 0.0323 | |
| | HIE+MSC_IN | 24 | 28.25 | 0.64 | | | 0.0002 | 0.9934 | 0.0633 |
| 4B | Control | 24 | 633 | 60.1 | F (3, 92) = 18.93 | < 0.0001 | | | |
| | HIE | 24 | 1320 | 69.7 | | | < 0.0001 | | |
| | HIE+MSC_IV | 24 | 1071 | 81.8 | | | 0.0006 | 0.1047 | |
| | HIE+MSC_IN | 24 | 665 | 90.1 | | | 0.9912 | < 0.0001 | 0.0016 |
| 4C | Control | 23 | 73.3 | 5.7 | F (3, 88) = 75.43 | < 0.0001 | | | |
| | HIE | 23 | 14.2 | 1.7 | | | < 0.0001 | | |
| | HIE+MSC_IV | 23 | 10.9 | 1.1 | | | < 0.0001 | 0.8973 | |
| | HIE+MSC_IN | 23 | 32.1 | 2.6 | | | < 0.0001 | 0.0013 | 0.0001 |
| 4E | Control | 27 | 73.60 | 2.4 | F (3, 107) = 445.7 | <0.0001 | | | |
| | HIE | 28 | 13.83 | 0.76 | | | < 0.0001 | | |
| | HIE+MSC_IV | 28 | 10.96 | 0.50 | | | < 0.0001 | 0.4400 | |
| | HIE+MSC_IN | 28 | 32.35 | 1.04 | | | < 0.0001 | < 0.0001 | < 0.0001 |
| 5B | Control | 6 | 38.25 | 5.86 | F (3, 20) = 13.99 | < 0.0001 | | | |
| | HIE | 6 | 15.07 | 3.28 | | | 0.0020 | | |
| | HIE+MSC_IV | 6 | 8.48 | 0.63 | | | 0.0001 | 0.6289 | |
| | HIE+MSC_IN | 6 | 33.94 | 3.73 | | | 0.8571 | 0.0120 | 0.0008 |
| 5C | Control | 6 | 44.02 | 2.48 | F (3, 20) = 12.69 | < 0.0001 | | | |
| | HIE | 6 | 28.70 | 2.32 | | | 0.0029 | | |
| | HIE+MSC_IV | 6 | 24.32 | 1.87 | | | 0.0002 | 0.6517 | |
| | HIE+MSC_IN | 6 | 40.75 | 3.61 | | | 0.8190 | 0.0207 | 0.0015 |
| 5D | Control | 6 | 0.087 | 0.011 | F (3, 19) = 6.351 | 0.0036 | | | |
| | HIE | 6 | 0.177 | 0.037 | | | 0.0262 | | |
| | HIE+MSC_IV | 5 | 0.143 | 0.008 | | | 0.2717 | 0.6903 | |
| | HIE+MSC_IN | 6 | 0.064 | 0.006 | | | 0.8541 | 0.0046 | 0.0709 |
